# Probiotic wheat sprouts: A novel functional food developed through *Lacticaseibacillus casei* inoculation with improved bioactivity and probiotic survival

**DOI:** 10.64898/2026.06.01.729244

**Authors:** Hasan Hajjami Barkousaraei, Mojtaba Mohammadzadeh Vazifeh, Mohammad Yaghoubi-Avini, Golnaz Shambayati

**Affiliations:** Faculty of Modern Biological Sciences and Technologies, University of Science and Culture, Tehran, Iran; Department of Microbiology, University College of Allameh Helli, Chalus, Iran; Faculty of Life Sciences and Biotechnology, Shahid Beheshti University, Tehran, Iran

**Keywords:** Wheat sprout, Probiotic, Non-dairy probiotic, *Lacticaseibacillus casei*, Bacterial inoculation

## Abstract

In this research we inoculated *Lacticaseibacillus casei* (*L. casei*) into wheat sprouts and studied the viability of the bacteria in the sprout.

*L. casei* (ATCC39392) strain was inoculated into sterilized wheat sprouts. The height of the sprouts and roots were checked and the active phenolic content, flavonoid compounds, and the antioxidant activity were measured. The bacterial viability was determined under simulated gastrointestinal (SGI) conditions. The physicochemical properties of the final product and its organoleptic properties were also investigated. The final confirmation of the presence of bacteria was also done by transmission electron microscope imaging.

The number of bacteria increased from 8.18 Log CFU/g to 11.81 ± 0.33 Log CFU/g. The increase in the phenolic content and antioxidant activity indicates the improvement of the nutritional value of the sprout. The physicochemical properties of the product changed due to the activity of bacteria. The inoculated bacteria also survived after exposure to SGI. The organoleptic properties of the product did not reveal a significant difference between the control and treatment groups.

The increase in the number of bacteria and their survival after exposure to SGI indicates the suitable condition of wheat sprout as a proper substrate for *L. casei* bacteria.

## 1. Introduction

The idea that food can act as medicine was first proposed by Hippocrates, who once said: “Let food be thy medicine and medicine be thy food.” (Soccol et al., 2010) A healthy diet is defined as a diet that includes the consumption of fruits, vegetables, fish, whole grains, and other foods that are beneficial for well-being (Wu et al., 2019). Over the past twenty years, quality of life has been recognized as an essential factor in the prevention and treatment of many diseases. The components and quality of a person’s diet are associated with five of the ten leading causes of death in the United States, including coronary heart disease, non-insulin-dependent diabetes mellitus, atherosclerosis, stroke, and various cancers (Corle et al., 2001).

Food security is a major concern in today’s societies, with an estimated one-eighth of the world’s population lacking access to adequate food (Daniels Jr & Morton, 2023). It is defined as the availability of sufficient food for all people at all times for an active and healthy life, which has two main characteristics: first, safe and nutritionally adequate food is available to all, and second, everyone should have access to acceptable food through socially acceptable means (i.e., not having to steal food from others to access it) (Anderson, 1990). Not having a healthy, balanced diet can also harm the body’s microbiome. Research has shown that disruptions in the human microbiome can contribute to a wide range of diseases, and there seems to be a close relationship between the state of the microbiome, health, and diet, suggesting that health enhancement can be modulated through diet and subsequent changes in the microbiota (Martinez et al., 2021).

Functional foods are food products that are used to improve health. The International Life Sciences Institute (ILSI) in North America defines functional foods as: “foods that provide health benefits beyond their primary nutritional value through physiologically active food components.” (Milner, 2002) Functional ingredients include polyunsaturated fatty acids (PUFA), antioxidants, probiotics, prebiotics, and synbiotics (Granato et al., 2010).

The word probiotic comes from the Greek word meaning “for life” (Kandylis et al., 2016). Probiotics are defined as live microorganisms that, when administered in adequate amounts, confer health benefits to the host (Granato et al., 2020). These microorganisms play a crucial role in enhancing digestive tract activity and promoting health through various mechanisms, including increased secretion of organic acids and the subsequent reduction in pH, as well as the production of bacteriocins (Sanders et al., 2018). Their consumption is also effective in improving or preventing many diseases, both physical and mental (Žučko et al., 2020). The primary benefit of probiotics in the industry is their application in dairy product production (Gao et al., 2021). However, today, due to factors such as increasing demand for vegetarianism, lactose intolerance, and allergies to milk cholesterol, the production of non-dairy probiotic products has received greater attention (Silanikove et al., 2015).

*Triticum aestivum*, also known as bread wheat, has been a primary source of human food and energy for centuries and is credited with being one of the main reasons for the success of the Roman Empire, earning it the nickname “the wheat empire.” (Igrejas & Branlard, 2020) It is estimated that today, about 55% of the carbohydrates consumed by humans come from wheat (Kumar et al., 2011).

Sprouting is a proven method for improving the nutritional properties of grains (Waliat et al., 2023). The sprouting process can be considered as a pre-digestion process during which the number of compounds with higher nutritional value increases (Marton et al., 2010). Wheat sprouts have higher antioxidant properties and a higher amount of free and bound phenolic compounds than unsprouted ones (Žilić et al., 2014).

The main purpose of this research was to produce a beneficial probiotic product by inoculating the probiotic bacteria *Lacticaseibacillus casei (L. casei)* 39392 into wheat sprouts and evaluating its nutritional value.

## 2. Materials and methods

### 2.1 Materials

The probiotic strain *L. casei* ATCC 39392 was obtained from the Iberesco Collection (Iberesco Life Science Co., Iran). MRS Agar, MRS Broth, and Plate Count Agar (PCA) (Iberesco Life Science Co., Iran) were used as microbial culture media in this study. Sodium hypochlorite, sodium hydroxide, hydrochloric acid, and a Gram staining kit were obtained from Ghatran Shimi (Ghatran Shimi Tajhiz Co., Iran). Pepsin and pancreatin powder were also manufactured by Sigma-Aldrich (Sigma-Aldrich Co., St. Louis, MO, USA).

### 2.2 Probiotic bacteria revival

MRS broth culture medium was sterilized by autoclaving at 121°C for 15 minutes. *L. casei* was cultured in a sterilized MRS broth medium and incubated at 37°C for 24 hours (Kusmiyati et al., 2020; Jung et al., 2021).

### 2.3 Production of probiotic wheat sprouts

#### 2.3.1 Seed surface disinfection

The seeds were soaked in 70% ethanol for 3 minutes after initial washing with sterilized distilled water, and then in 3.5% hypochlorite for 30 minutes after washing with sterilized distilled water to prevent the growth of lactobacilli. The seeds were then washed three times with sterilized distilled water and transferred to a sterile plate (Tverdokhlib et al., 2018).

#### 2.3.2 Seeds germination

The disinfected wheat seeds that were visibly damaged during the disinfection process were separated, and the remaining seeds were incubated at 25°C for 24-48 hours to induce germination (Banerjee & Mittra, 2018).

#### 2.3.3 Inoculation of probiotic bacteria into sprouts

The MRS broth medium containing *L. casei* strain 39392 was separated using a Rotofix A32 centrifuge (Hettich Instruments, Tuttlingen, Germany) at 4500 rpm for 10 minutes at room temperature, after washing twice with Phosphate-buffered saline (PBS) (Merck, Kenilworth, NJ, USA), and the bacterial precipitate was collected (Son et al., 2018). The treatment sprouts were soaked in a suspension containing 1.5e+8 CFU/ml bacteria (0.5 McFarland) for 60 minutes. The control group was also soaked in a PBS solution (containing no bacteria) for 60 minutes and then removed. All the sprouts were then separated from the suspensions (Tverdokhlib et al., 2018).

#### 2.3.4 Sprouts growth

In separate plates, a sterilized filter paper was moistened with autoclaved tap water at 121°C for 15 minutes, and then 6 grams of sprouts were placed in each plate. The samples were divided into a control group without inoculation and a treatment group inoculated with probiotic bacteria. The plates were incubated for 5 days at 25°C and then harvested for further analysis (Egamberdieva, 2009; Žilić et al., 2014).

### 2.4 Examination of the characteristics of sprout growth

Growth characteristics, including root height and size of sprouts, were measured using a caliper on day 5 of inoculation (Egamberdieva, 2009; Limanska et al., 2018).

### 2.5 Probiotic count of sprout

1 g of treated sprout was washed with a vegetable disinfectant solution (Man Co, Iran) and sterile distilled water, and then a serial dilution was prepared. The probiotic (*L. casei*) count was done by the pour plate method with MRS agar medium on days 0 and 5 after inoculation (Vazifeh et al., 2021; Ortolani et al., 2007).

### 2.6 Measurement of total flavonoid, total phenolic compounds, and antioxidant activities in sprouts

After cleaning and washing, the samples were transferred to an oven at 40°C until their moisture content reached about 10%. Then, the samples were ground and prepared. For extraction, 2 g of powdered sprout tissue was added to 20 ml of pure methanol and shaken for 72 hours. The samples were then ultrasonicated for 30 minutes and finally centrifuged at 6000 rpm for 10 minutes. The upper extract was used to measure phenol, flavonoid, and antioxidant properties (Esmailzadeh et al., 2023).

Total flavonoids were measured by the aluminum chloride colorimetric method and by a spectrophotometer at an absorbance of 425 nm (Pękal & Pyrzynska, 2014). The results were expressed as micrograms per milliliter of extract. Total phenol content was measured using the Folin-Ciocalteu colorimetric method. The results were expressed as equivalent values of gallic acid concentration at an absorbance of 760 nm (Bancuta et al., 2016).

The antioxidant properties of the sprout extract were investigated for 15 minutes at room temperature in the dark using 2,2-diphenyl-1-picryl-hydrazyl (DPPH). Then, the absorbance of the mixture was read at 517 nm and calculated based on the absorbance of methanol as a blank sample according to the following formula:

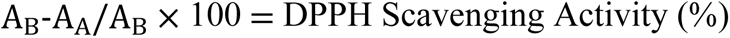

where A_B_ is the absorbance of the blank, and A_A_ is the absorbance of the sample (Padmanabhan & Jangle, 2012; Okawa et al., 2001).

These tests were performed by the Research Institute of Forests and Rangelands (Tehran, Iran) for more accurate results.

### 2.7 Measurements of physicochemical properties

#### 2.7.1 pH

The pH of both control and treatment samples was measured using a pH meter. After calibrating the device, 1 g of the sample was mixed with 50 ml of distilled water, and the total volume was adjusted to 100 ml. The pH was then read twice, at the first minute with stirring and the second minute without stirring (ISO 8700, 2025; OECD, 2013).

#### 2.7.2 Acidity

To determine acidity, 5 g of each sample was titrated with 0.1N sodium hydroxide. The acidity of both samples was measured using the following formula, and calculated as the degree of oleic acid, which is the main fatty acid in a wheat sprout (Nemzer & Al-Taher, 2023).

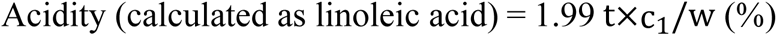

where t is the normality (mol/L) of NaOH, c is the volume (ml) of NaOH, and w is the weight (g) of the sample (OECD, 2013; AOCS, 2017; Shambayati et al., 2025)

### 2.8 Dry matter measurement

Dry matter was determined by drying the weighed samples in an oven at 78℃ for 24h. The following formula was used to measure the percentage of dry matter (Erkovan et al., 2008):

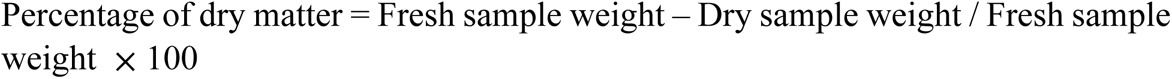

### 2.9 Viability of bacteria under simulated gastrointestinal conditions

#### 2.9.1 Examination of the viability of bacteria under simulated gastric conditions

To prepare the simulated gastric juice, 2 grams of sodium chloride and 2.3 grams of pepsin powder were dissolved in distilled water, and the final volume was made up to 1000 ml. The pH of the medium was adjusted to 2.3 using hydrochloric acid. The final solution was autoclaved at 121°C for 30 minutes. 1 g of the sprout was then mixed with 9 ml of pepsin solution until as homogeneous as possible, and the mixture was incubated at 37°C for 3 hours. After incubation, dilutions were prepared from the mixture, plated on MRS agar using the pour plate method, and incubated at 37°C for 72 hours. The number of probiotic bacteria was counted before and after this test (Bao et al., 2010).

#### 2.9.2 Examination of the viability of bacteria under simulated intestinal juice

To simulate intestinal juice, pancreatin was suspended in sterile saline at a final concentration of 1 g/L, supplemented with 4.5% bile salts, and the pH was adjusted to 8.0 using sterile 0.1 mol/L NaOH (Blaiotta et al., 2013). 1 g of the sprout was mixed in 9 ml of the final solution as homogeneously as possible and incubated at 37 °C for 3 hours. The mixture was cultured in MRS agar medium using the pour plate method and incubated at 37°C for 72 hours. Probiotic bacteria were counted before and after the test (Bao et al., 2010).

### 2.10 Transmission electron microscope (TEM) imaging

To observe the presence of bacteria in the sample, after initial confirmation of its presence in transverse and longitudinal sections prepared from the treated sprouts using Gram staining, the samples were placed in fixative and sent to the transmission electron microscopy imaging unit at Pasteur Institute (Tehran, Iran) (Micci et al., 2022; Liu et al., 2022).

### 2.11 Organoleptic evaluation

Based on the hedonic sensory evaluation method, a table with 5 evaluable parameters, including color, aroma, taste, appearance, and texture, was prepared. 5 panelists were selected, and testing was carried out by giving a questionnaire containing 1-5 points (from the lowest to the highest) for each parameter to the panelists (AdebayoTayo & Akpeji, 2016).

### 2.12 Statistical analysis

The results of all experiments in this study were based on the average of at least three repetitions and were statistically analyzed using IBM SPSS Statistics 27 software and the T-test analysis (Vermeulen et al., 2008). Differences were considered significant at *P* ≤ 0.05.

## 3. Results and discussion

### 3.1 Comparison of growth characteristics in the treatment and control of wheat sprouts

After 5 days of inoculation, 10 samples were randomly selected from the treatment and control groups, and their characteristics (sprout and root height) were measured. The results indicate a significant (*P* ≤ 0.05) difference between the treatment and control groups, showing better growth in the treatment group (Fig. 1, 2).

**Figure 1.**
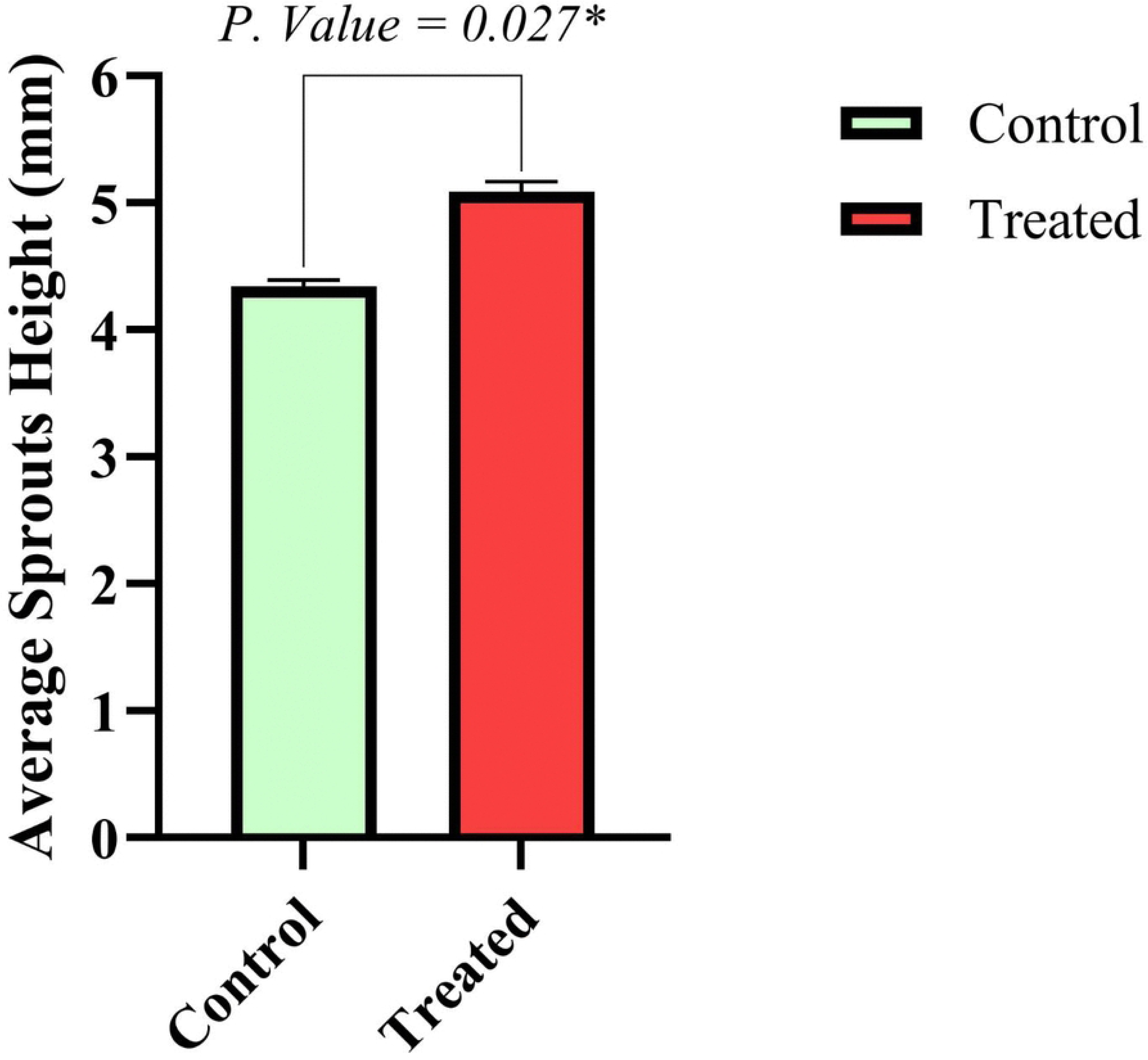
Average sprout height comparison of the probiotic and control samples

**Figure 2.**
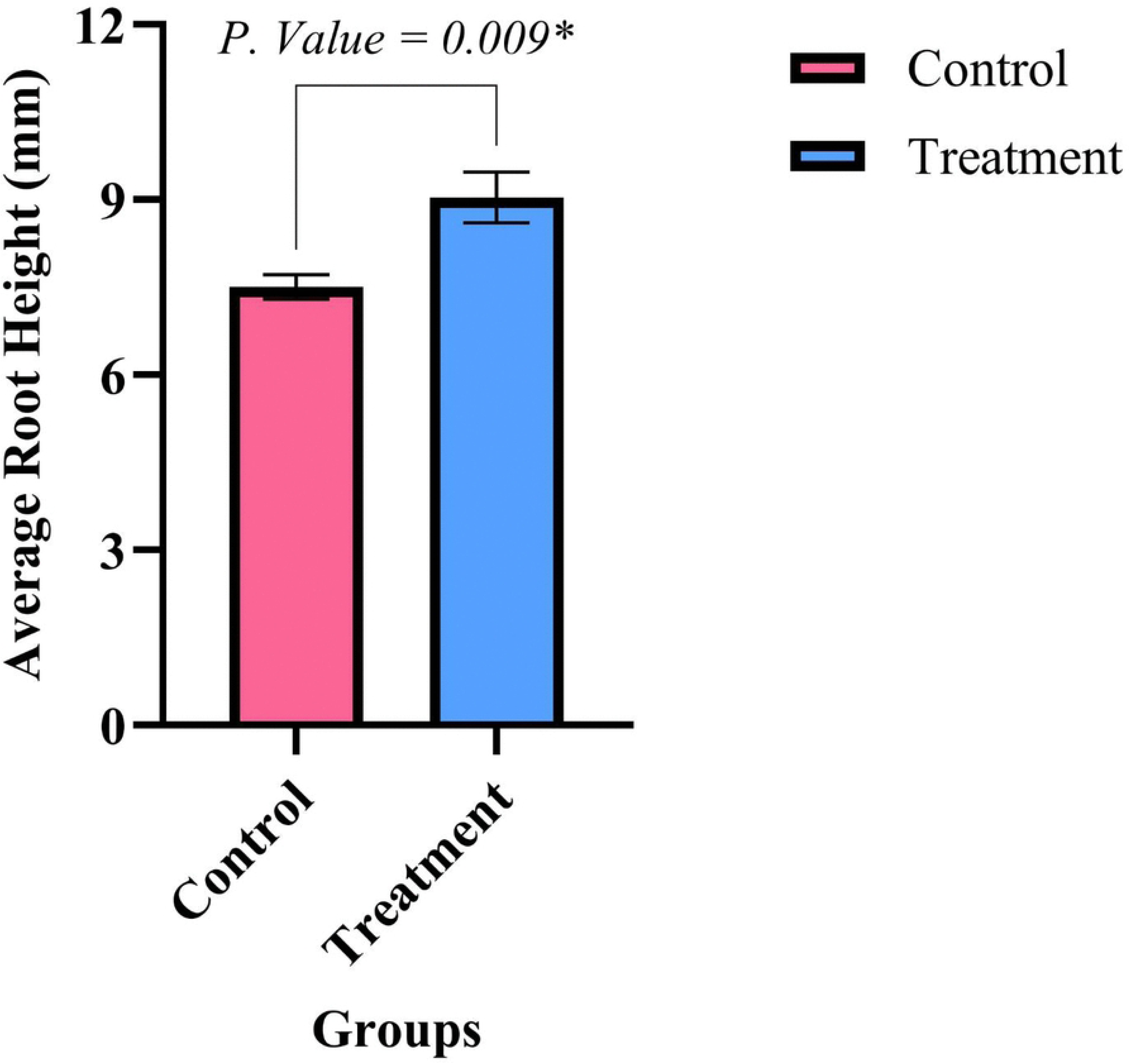
Average root height comparison of the probiotic and control samples

The significant increase in root length of treated sprouts compared to the control was 1.205 times on average, and the significant increase in sprout height was 1.169 times.

### 3.2 Probiotic count

No growth was observed in the sprout count before inoculation on MRS agar medium. The control germinated wheat after inoculation on day 0 and day 5, and the cultures did not show any growth on MRS agar medium. The count of probiotic sprouts on MRS agar medium after inoculation was close to the inoculum count, with a slight difference of 7.92 ± 0.09 Log CFU/g.

The probiotic counts on day 5 of the treatment sample were 11.66 ± 0.48 Log CFU/g, indicating an increase in bacterial numbers in wheat sprouts. According to the results obtained by counting the treated sprouts and the requirement for at least 6 Log CFU/g or ml of bacteria in probiotic products, treated sprouts can be considered a non-dairy probiotic product.

### 3.3 Total phenols, total flavonoids, and antioxidant properties in treated and control sprouts

#### 3.3.1 Total phenol

Total phenol of the bacteria-containing sample was 74.189 ± 0.419 and the control was 69.385 ± 0.395. These results indicate a significant (*P* ≤ 0.05) increase in the total phenol content in the probiotic sample compared to the control one. Total phenol increased by 6.92% in wheat sprouts due to the activity of probiotic bacteria (Fig. 3).

**Figure 3.**
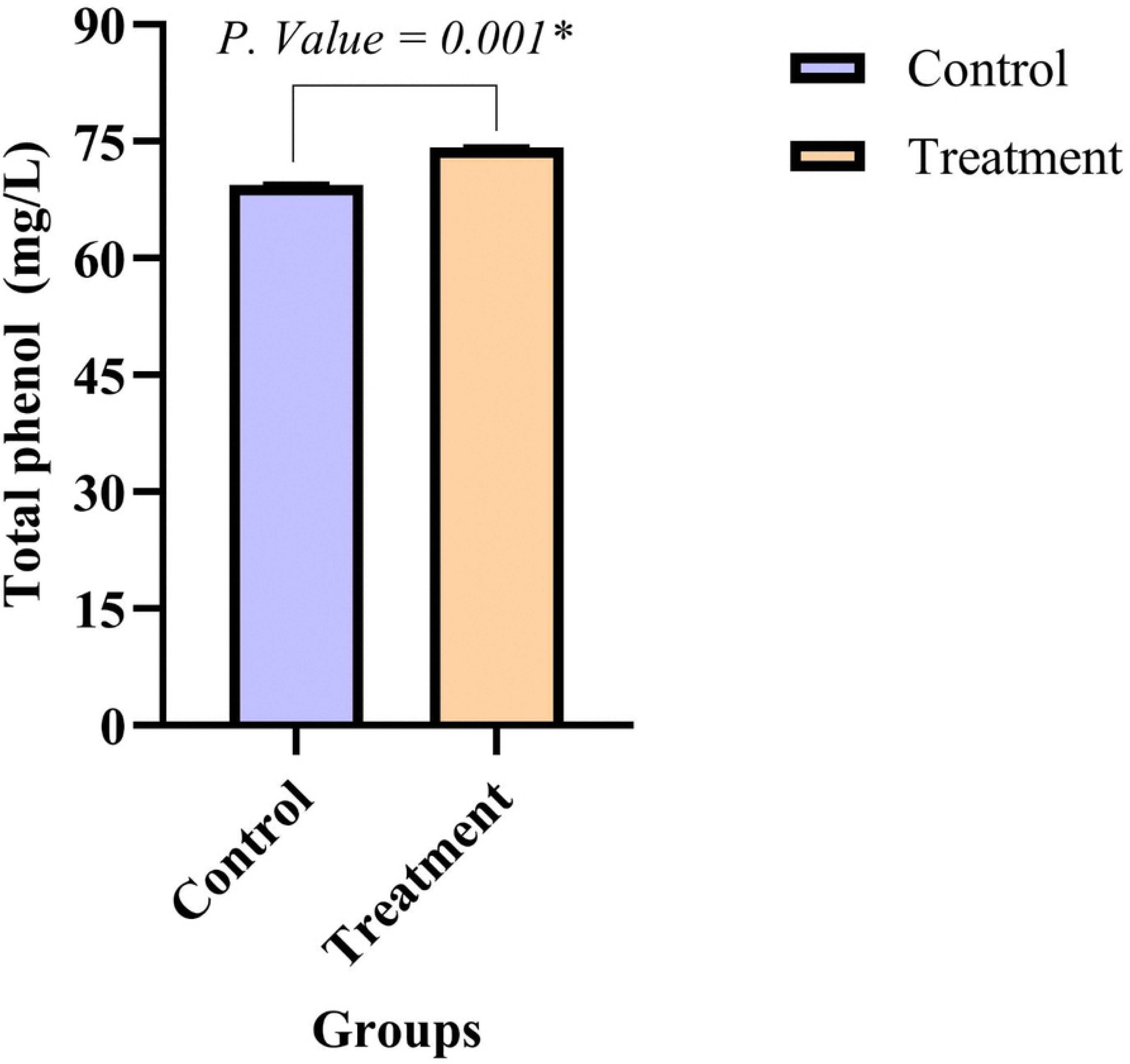
Total phenol content of the treatment and control samples

#### 3.3.2 Total flavonoids

Total flavonoid compounds in the probiotic sample were 12.073 ± 0.056 compared to the control group, which was equal to 10.95 ± 0.051 (Fig. 4) and increased significantly (*P* ≤ 0.05) by 10.24% compared to it.

**Figure 4.**
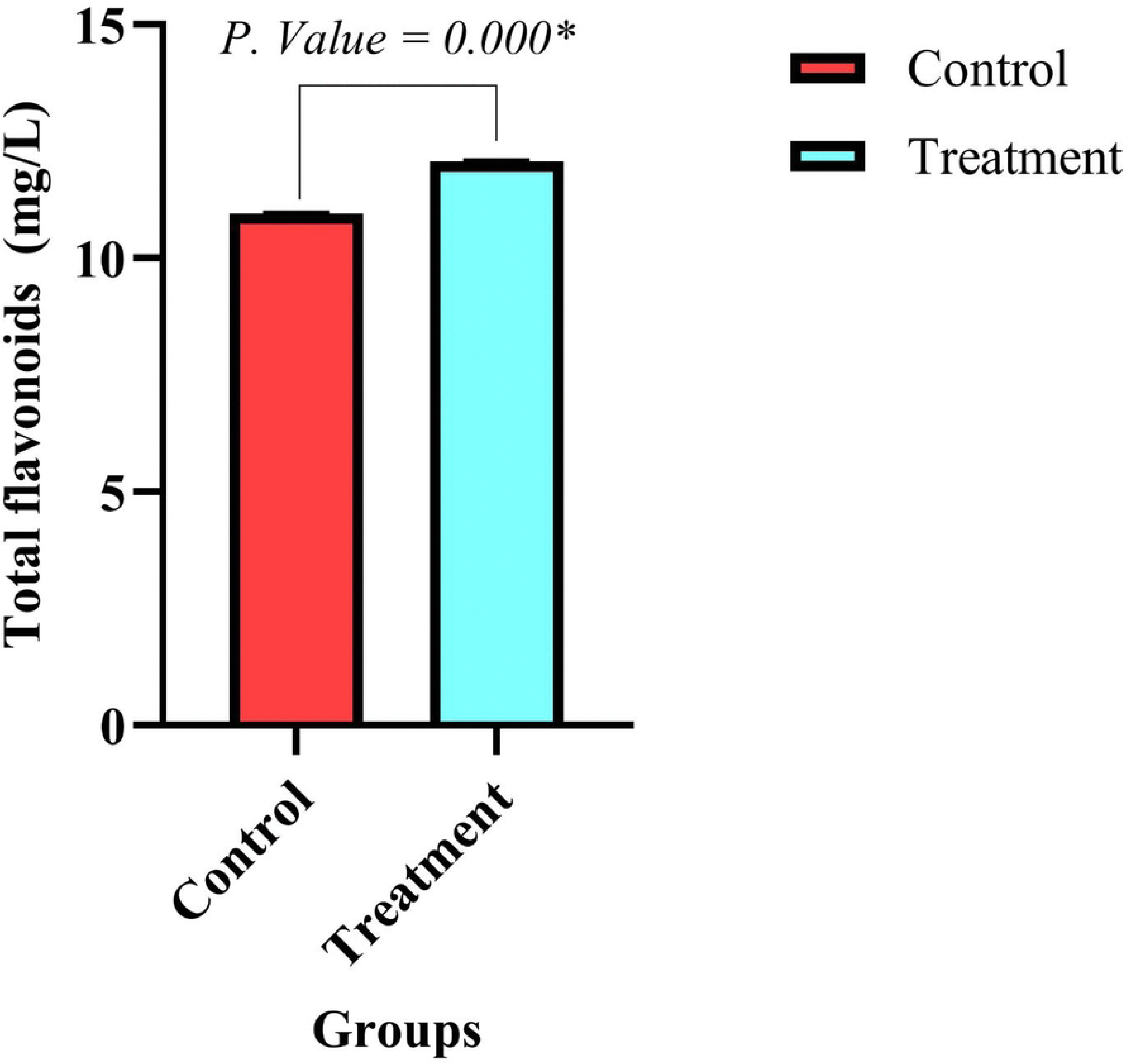
Total flavonoid content of the treatment and control samples

#### 3.3.3 Antioxidant properties

Antioxidant properties result in the control (54.83 ± 0.27) and treatment sample (56.5 ± 0.81) indicated a significant (*P* ≤ 0.05) difference (Fig. 5). The changes in antioxidant properties and their increase compared to the control is 2.955%.

**Figure 5.**
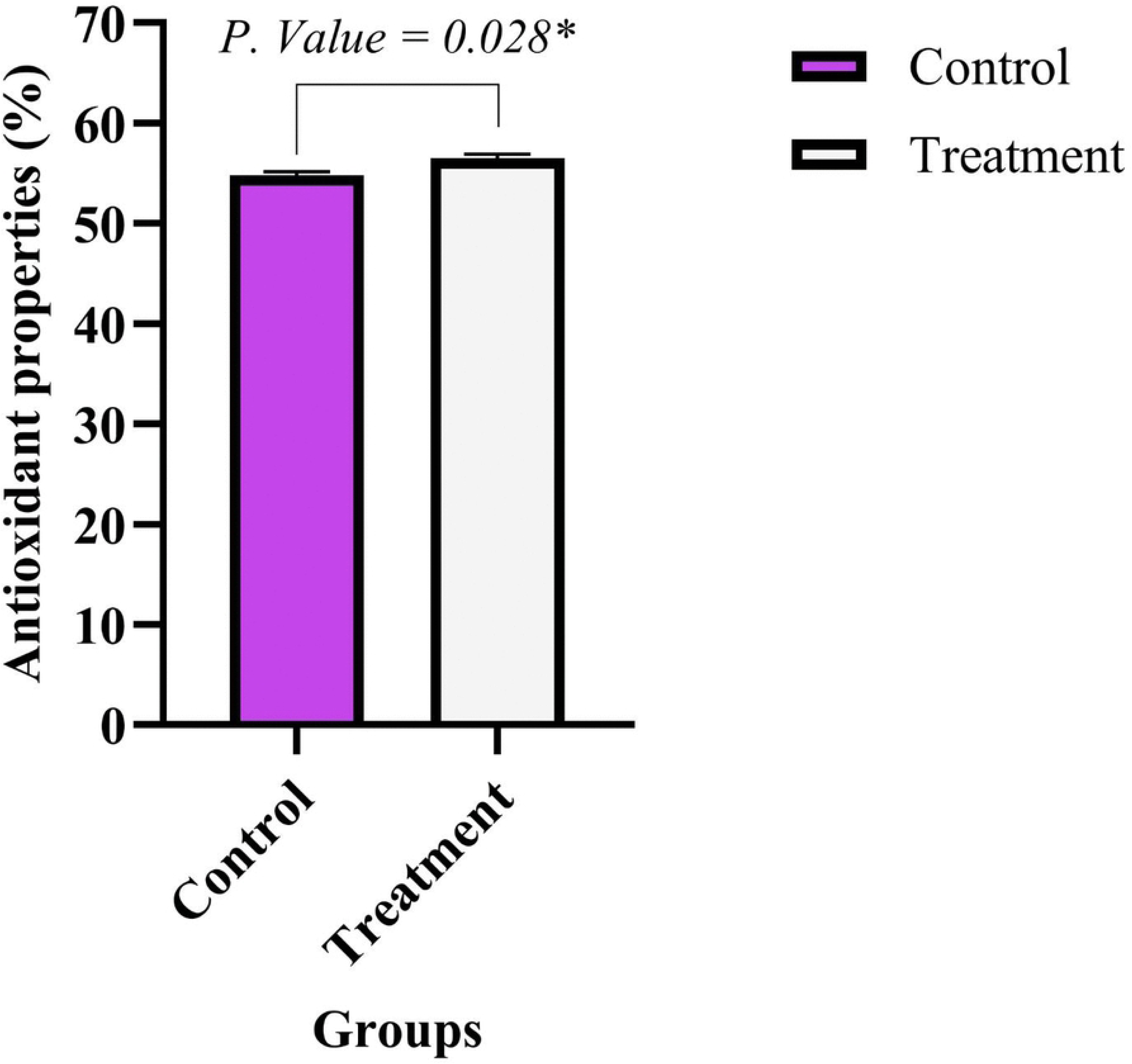
Antioxidant properties evaluation of the treatment and control samples

### 3.4 Measurement of physicochemical properties

#### 3.4.1 pH

The calculated pH values for the control and treatment samples at minute one were, respectively, 5.733 ± 0.321 and 4.783 ± 0.189 (Fig. 6a) and at minute two were equal to 5.87 ± 0.106 and 4.403 ± 0.242 (Fig. 6b). The average results indicate a significant (*P* ≤ 0.05) decrease in pH in the treatment sample compared to the control sample, which could be attributed to the acid production by *L. casei*.

**Figure 6.**
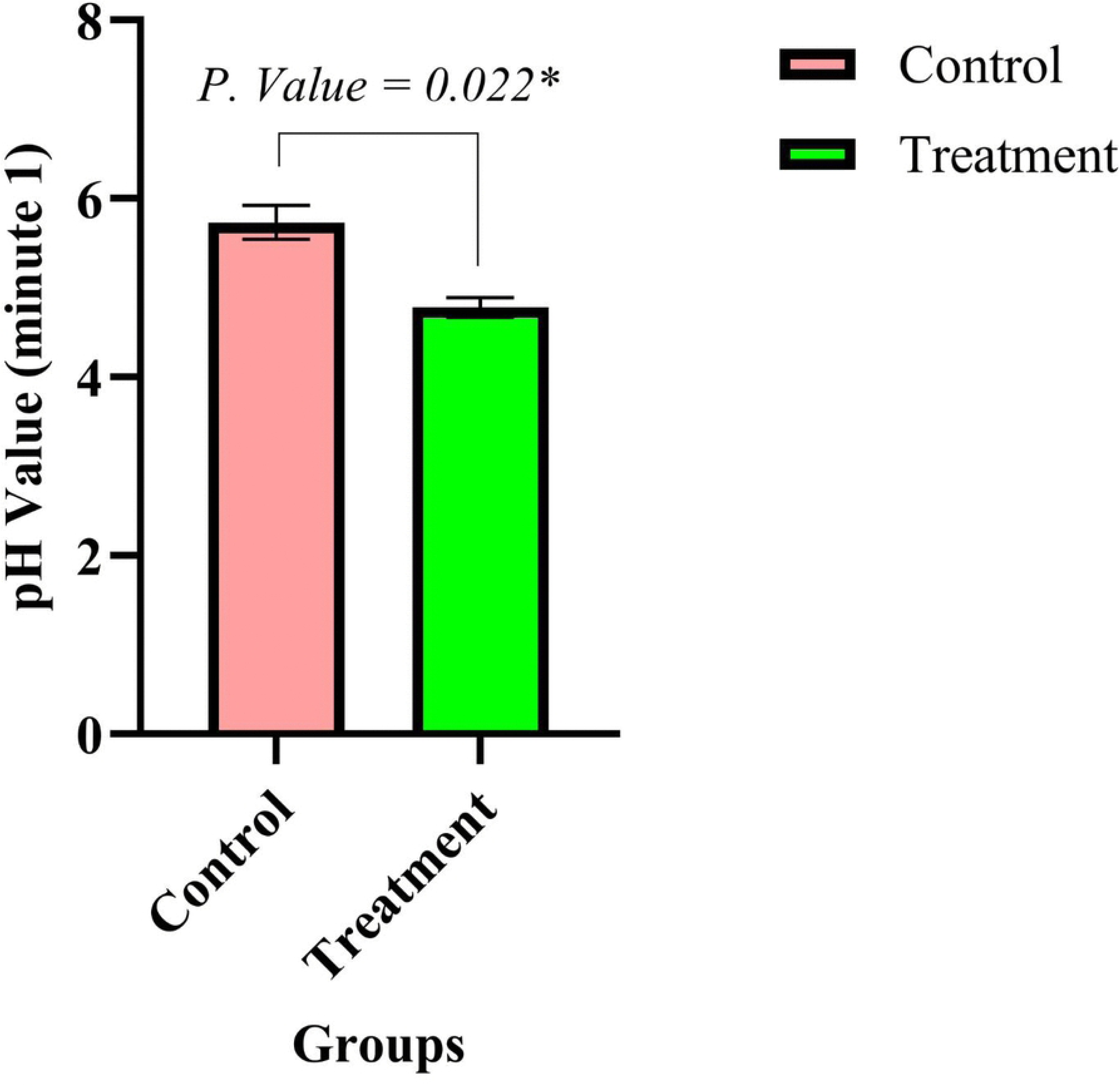
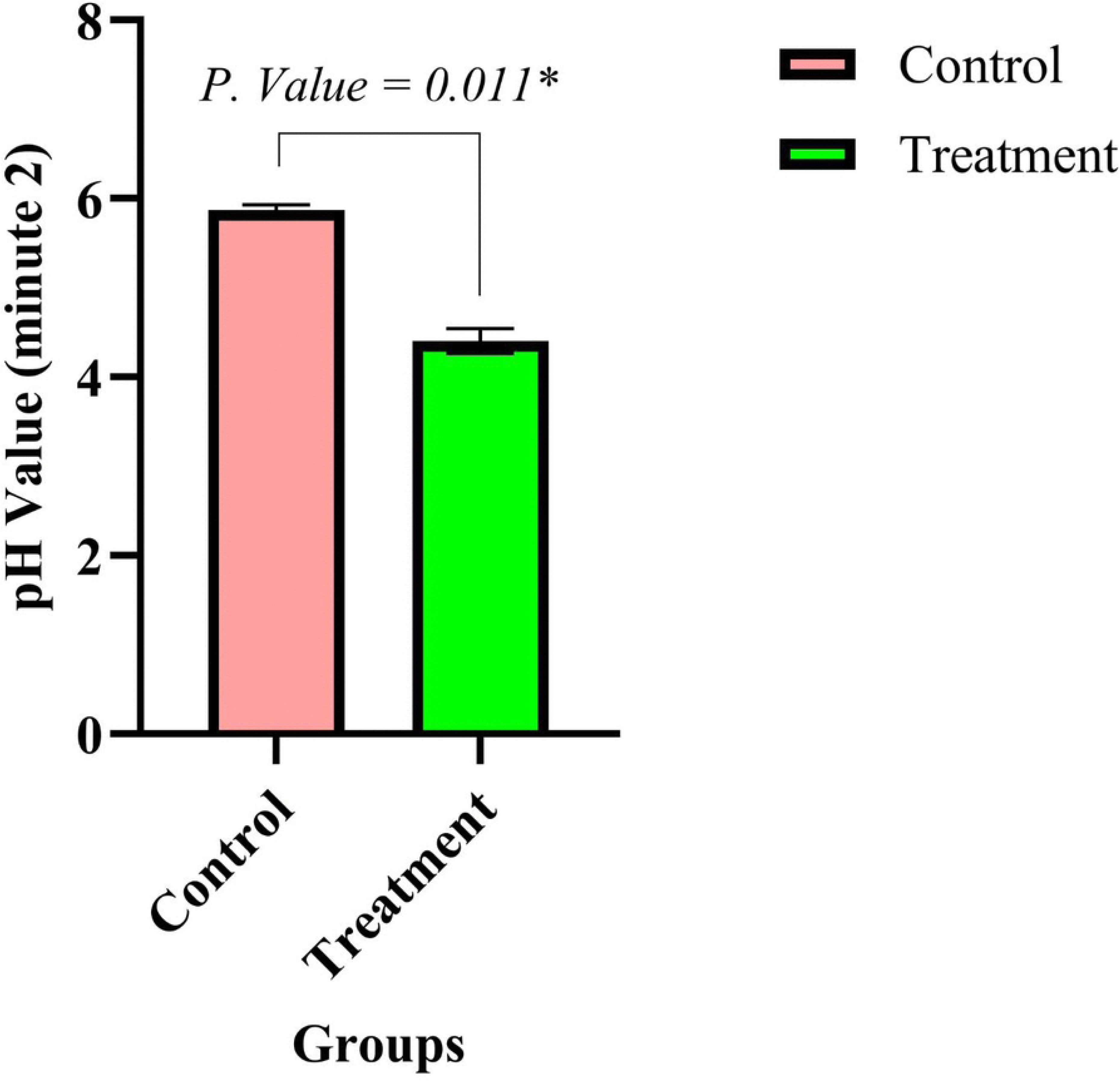
pH values of two samples at minutes 1 (a) and 2 (b)

#### 3.4.2 Acidity

The measured acidity in the treatment sample was equal to 0.017 ± 0.002, and in the control, it was 0.006 ± 0.001 (Fig. 7). These results indicated a significant (*P* ≤ 0.05) increase in acidity in the treatment compared to the control.

**Figure 7.**
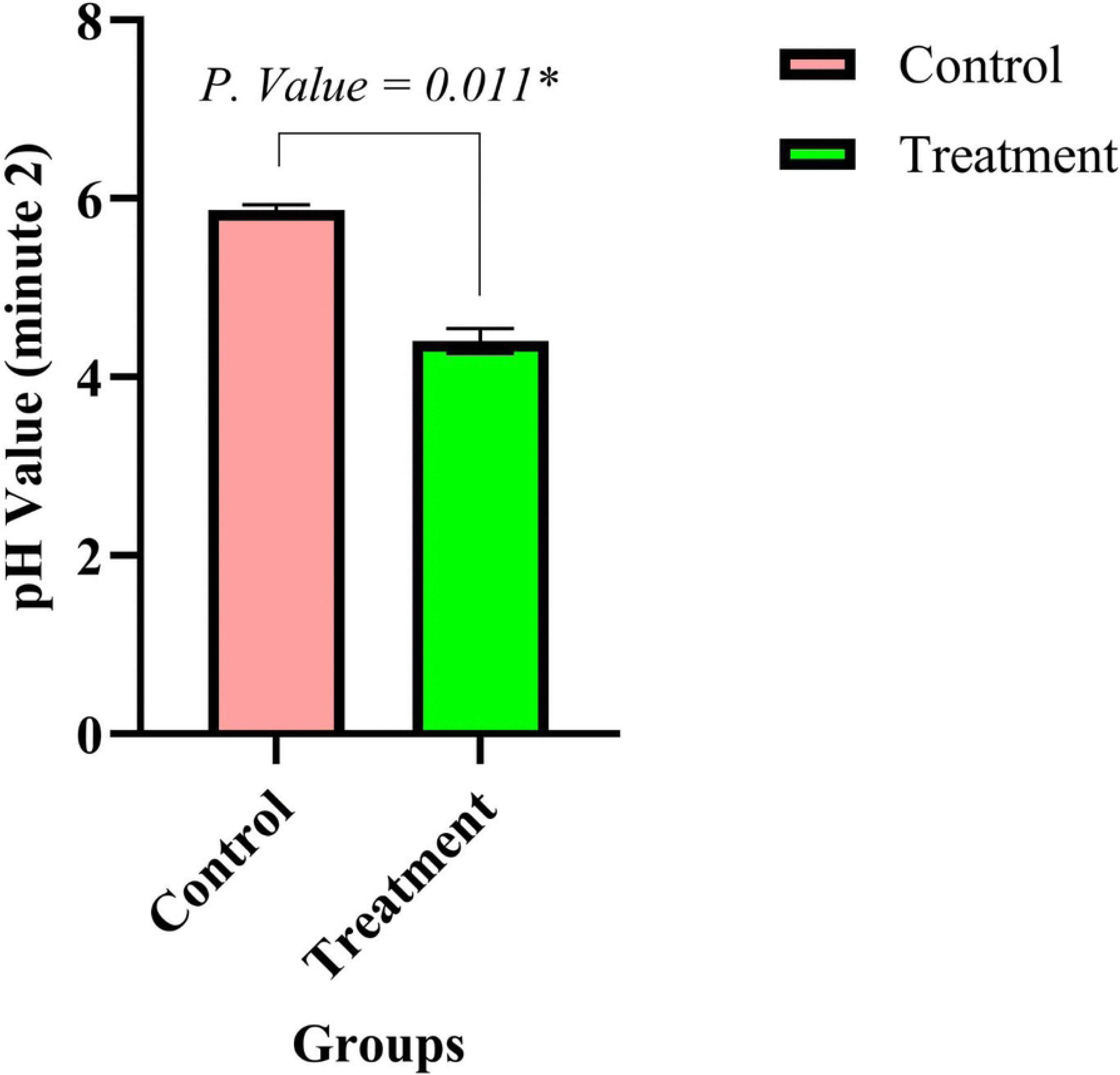
Acidity comparison of two samples, indicating higher values in the treatment sample

### 3.5 Dry matter

The dry matter content obtained for the treatment sample was 84.086 ± 0.05, and for the control sample was 83.980 ± 0.582, but no significant difference (*P* > 0.05) was observed in the analysis of the measurements related to the dry matter of the samples.

### 3.6 Gastrointestinal environment simulation

The number of probiotic bacteria inoculated into the sprout was counted before and after exposure to gastrointestinal juices (pepsin and pancreatin media). The results indicated that probiotic bacteria in the sprout were able to resist gastric juice. The number of bacteria before exposure to the simulated environment was 11.80 ± 0.18 Log CFU/g on average for three replicates, which decreased to 10.85 ± 0.13 Log CFU/g and 10.95 ± 0.04 Log CFU/g, respectively, after exposure to pepsin and pancreatin environments (Fig. 8, 9).

**Figure 8.**
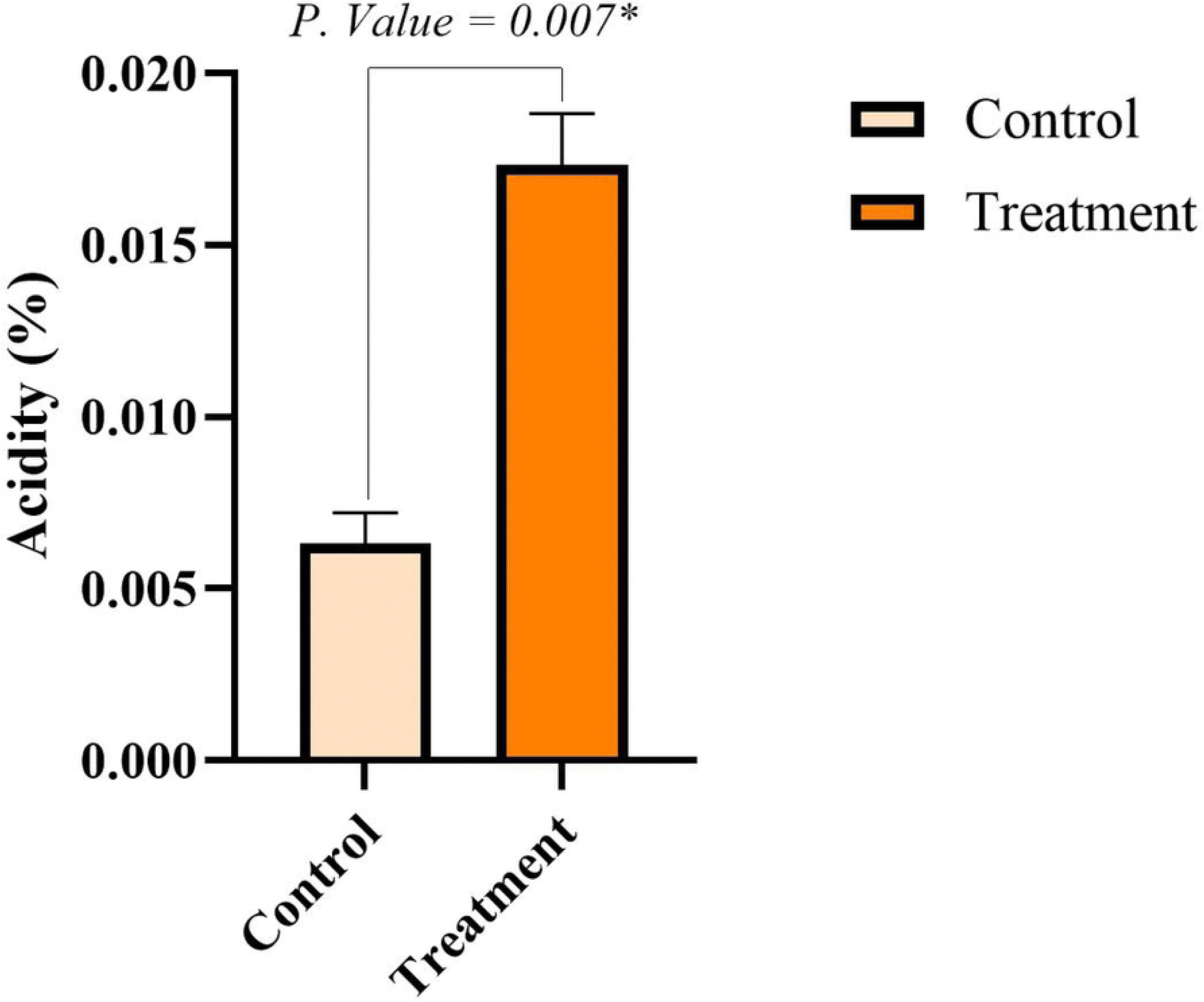
Colony count before and after exposure to pepsin

**Figure 9.**
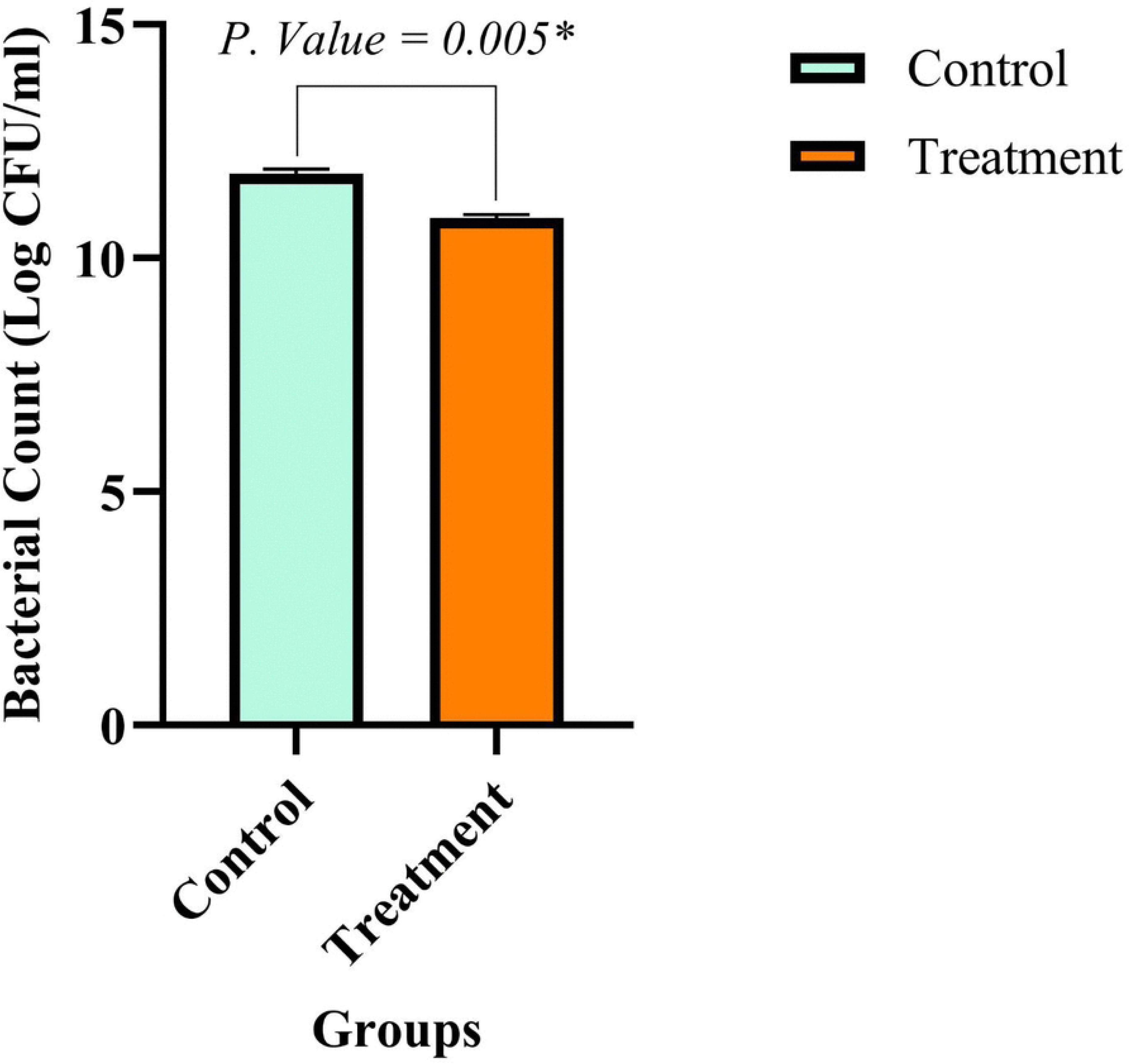
Colony count before and after exposure to pancreatin

### 3.7 Gram staining and TEM

Final confirmation of the presence of probiotic bacteria in the treated sample was obtained using TEM. The results of the initial Gram staining and TEM imaging are presented in Figures 10 and 11, respectively.

**Figure 10.**
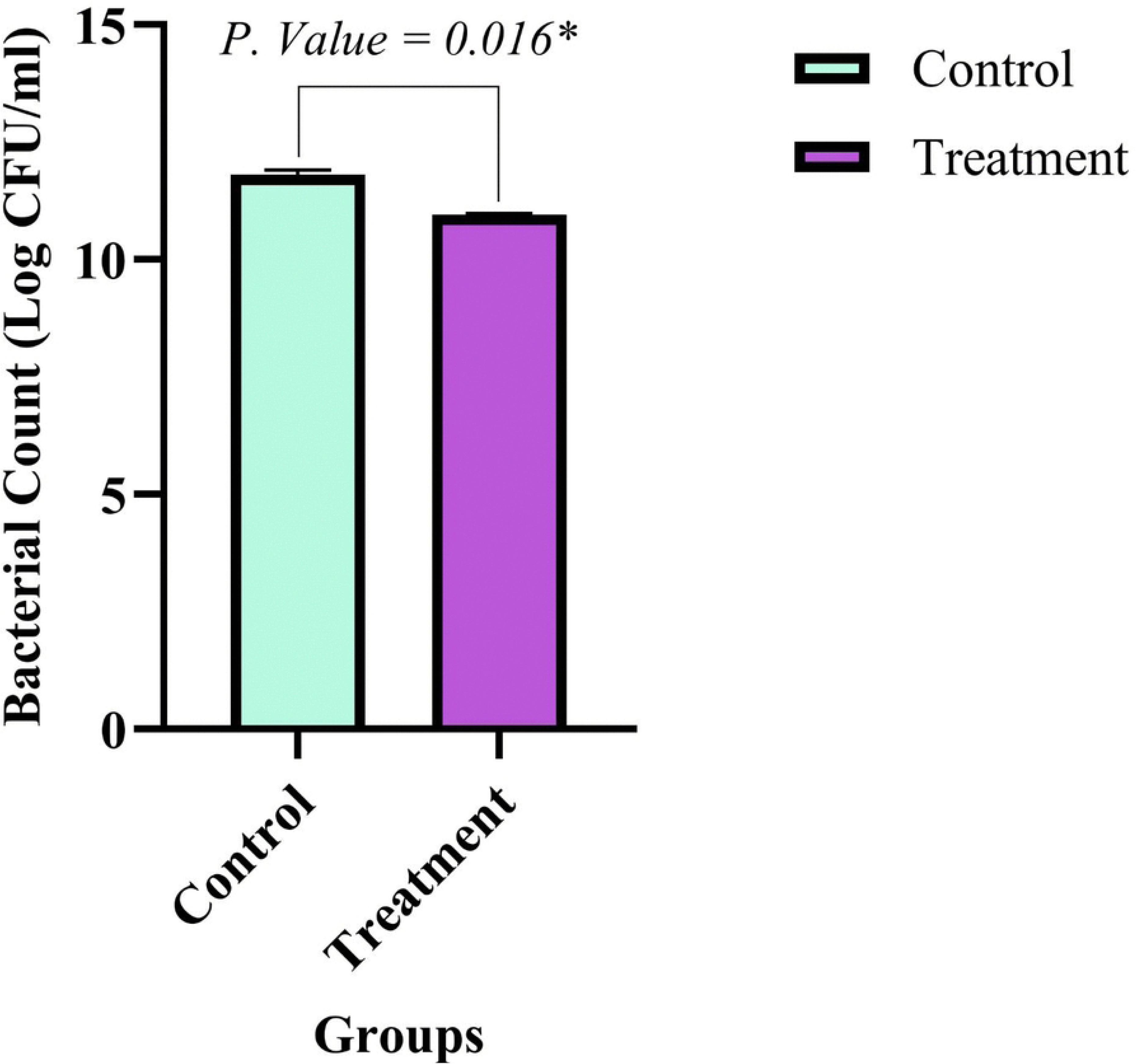
Gram staining of the transverse and longitudinal sections of the L. casei containing sample

**Figure 11.**
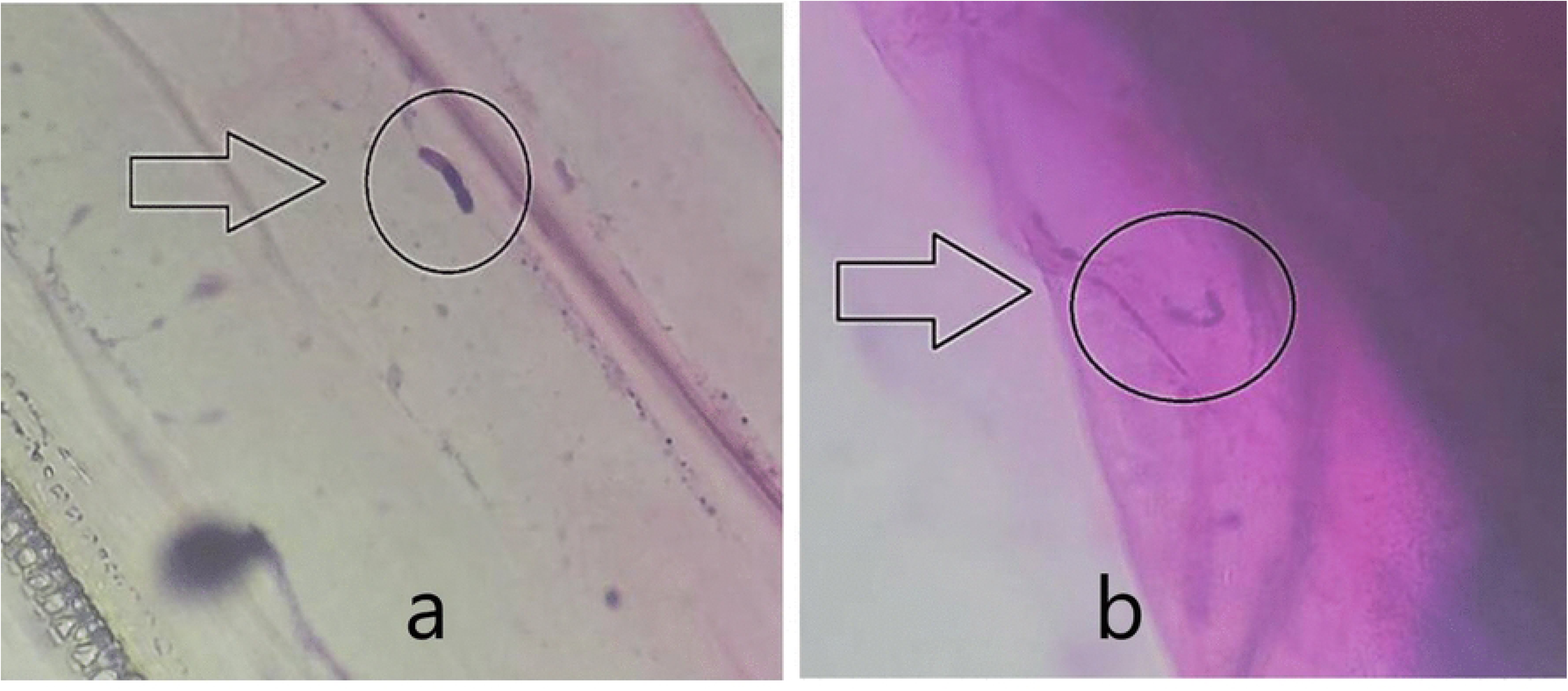
TEM imaging of L. casei-containing sample

### 3.8 Organoleptic evaluation

Although the texture and aroma of probiotic wheat sprouts were slightly higher, they were less acceptable in terms of appearance than the control sprout (Fig. 12). These results indicate that bacterial inoculation did not significantly (*P* > 0.05) affect the overall acceptability of the samples.

**Figure 12.**
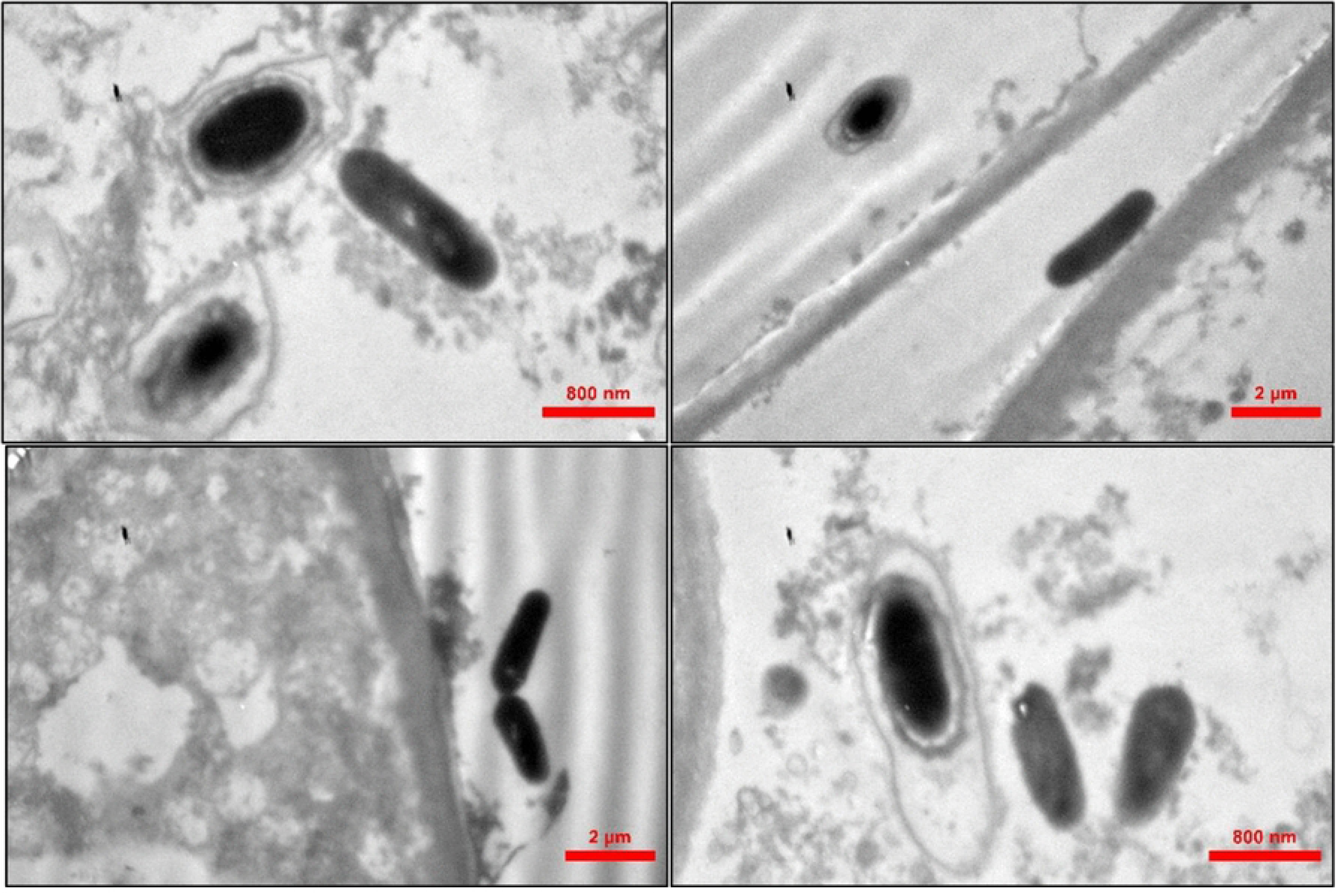
Organoleptic evaluation results are an average of fifteen determinations

**Figure 13.**
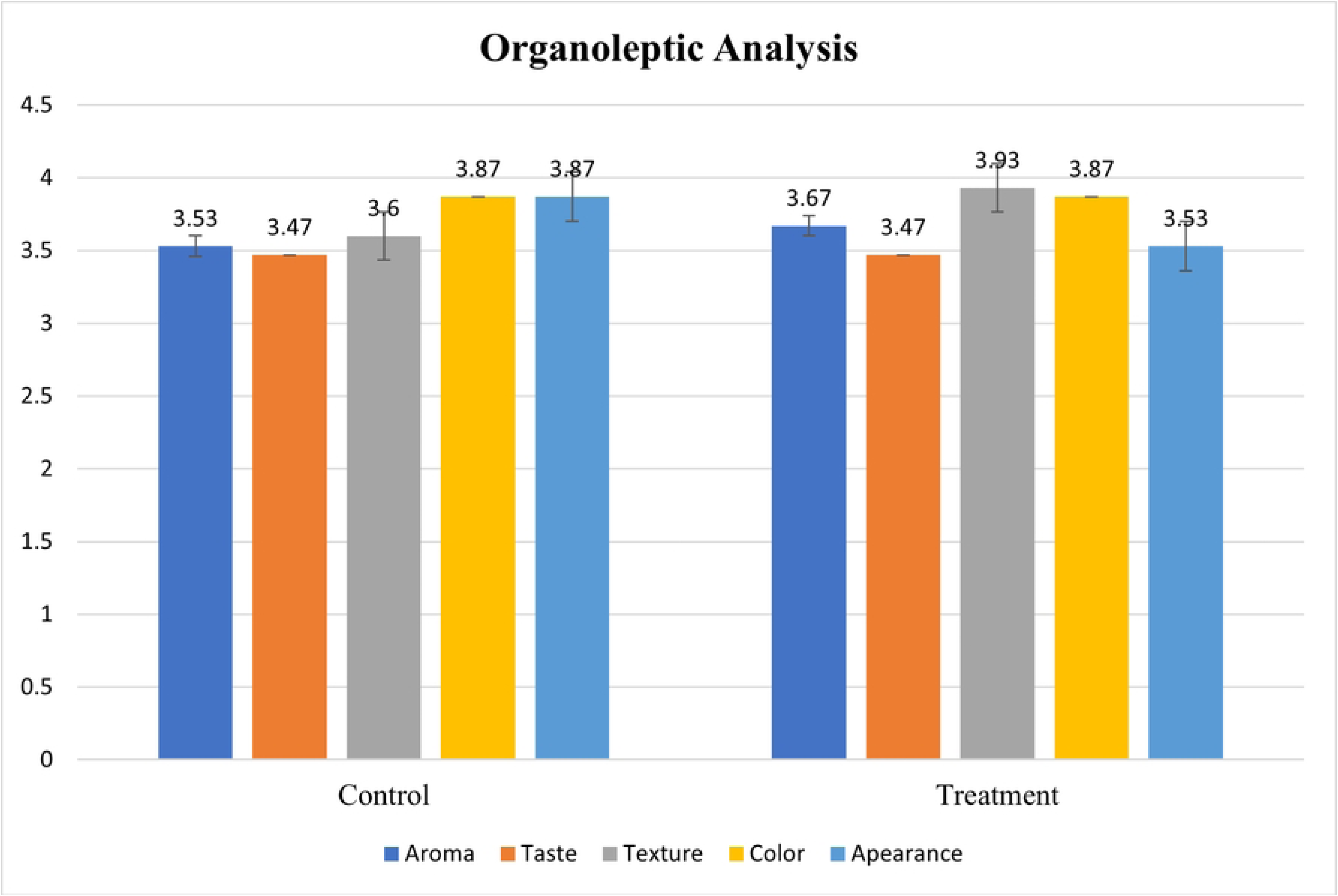

The results obtained regarding the effect of probiotic strain inoculation on root and sprout height are similar to the results of Limanska et al., who reported a 2.4-fold increase in root size and 1.228-fold increase in sprout height for treated samples compared to the control ones using *Lactobacillus plantarum* as a growth stimulant (Limanska et al., 2018). Tverdokhlib et al. also observed a 1.17-fold increase in root size and a 1.243-fold increase in sprout height for treated shoots when using a combination of several probiotic strains as a growth stimulant on wheat sprouts (Tverdokhlib et al., 2018).

Many studies have confirmed the effect of probiotic inoculation on the phenolic content and antioxidant properties of different products. In their study on the effects of fermentation of a type of red algae by *Lactobacillus brevis*, Wang et al. observed that the addition of this probiotic bacterium increased the total phenol content and increased the antioxidant properties of the final product (Wang et al., 2023). It is similar to the results of the study by Song et al. on the effect of using the probiotic bacteria, *Pediococcus acidilactici,* on the fermentation of black raspberry extract, in which they reported that the use of these probiotic bacteria, in addition to increasing the amount of phenols and total flavonoids, also played a role in increasing the antioxidant effect of this drink (Song et al., 2021). Holkem et al. obtained the same result using a combination of *Bifidobacterium animalis* and *Lactobacillus paracasei* bacteria on blueberry fruit extract (Holkem et al., 2021). Moreover, Shori et al. used three probiotic bacteria, *Lacticaseibacillus casei*, *Lactobacillus rhamnosus*, and *Lactobacillus plantarum*, in starter cultures separately during the study of cashew milk fermentation for yogurt production, and found that the antioxidant activity of the product, total phenols, and total flavonoids increased in all three treatments (Shori et al., 2022). The increase in antioxidant properties, total phenolic content, and total flavonoid content in the results mentioned above, due to the presence of probiotic bacteria, is similar to the results obtained in this study.

In confirmation of the obtained results in the enumeration and survival of bacteria and the decrease in the number of them during the storage period, Zheng et al. observed a slight decrease in the number of bacteria counted after 4 weeks of storage of a beverage after the addition of the probiotic strain *L. casei* to it (Zheng et al., 2014), which is similar to the results reported by Shambayati et al. in the production of probiotic garden cress (Shambayati et al., 2025). A decrease in the number of probiotic strains in different products, such as cashew apple juice, after 21 days of storage has been observed and reported (Pereira et al., 2011). Angelov et al., in a study on the production of a fermented beverage from whole barley using *Lactobacillus plantarum*, reported that the number of inoculated bacteria increased from 0.5 McFarland to 7.5 × 10^10^ CFU/ml over an 8-hour fermentation. The number of remaining bacteria also reached 10^7^ CFU/ml after 24 days of storage (Angelov et al., 2006). Champagne and Gardner investigated the viability of *Lactobacillus plantarum*, *Lactobacillus reuteri*, and *Lactobacillus rhamnosus* in a commercial juice and reported that the bacteria were able to survive in the drink for up to 80 days at 7 Log CFU/ml and that it is possible to deliver probiotics through products based on the type and composition of the selected substrate (Champagne & Gardner, 2008). A decrease in pH is usually observed in fermented products produced with probiotics (Sanders et al., 2018), but on the other hand, various studies have shown that the presence of probiotics does not affect the pH or titratable acidity of the product (Tripathi & Giri, 2014). Consistent with most studies, we found a lower pH and greater acidity in the sample containing Lactobacillus than in the control. Yoon et al., in the production of a probiotic fermented beverage using *L. plantarum*, observed a decrease in the pH of the final product from 6.3 to 1.4 and an increase in its acidity from 0.13 to 0.56, due to the activity of the aforementioned bacteria and acid production (Yoon et al., 2005). Teneva et al. reported that the pH of the mayonnaise using basil extract and *L. plantarum* decreased from 6.5 to 4.5 during a 20-day storage period due to the activity of the bacteria (Teneva et al., 2020). These results are like those of Sagdic et al. in the production of probiotic ice cream and its pH decrease and acidity increase during storage (Sagdic et al., 2012).

As in other reports, the dry matter content of the probiotic sample was not significantly higher than that of the control sample. Gedic et al. reported that in the production of a fermented dairy beverage using *L. casei*, *L. rhamnosus*, and *L. acidophilus*, the amount of dry matter in the final product did not differ significantly from that of the control group without probiotic bacteria (Gedik & Karahan, 2023). Sagdic et al. also reported that the dry matter content of probiotic ice cream produced with *L. casei* bacteria was not significantly different from that of non-probiotic ice cream (Sagdic et al., 2012). In contrast, Oliveira et al. observed an increase in the dry matter content of goat cheese and suggested that this might be linked to higher protein levels. Therefore, they concluded that similar changes in dry matter may occur in dairy probiotic products (Oliveira et al., 2012).

The survival of probiotic strains under simulated gastrointestinal (SGI) conditions is crucial for delivering their health benefits. A rapid loss in the number of free bacterial cells in SGI conditions is observed in many studies (Casarotti et al., 2015; Chávarri et al., 2010; Khorasani & Shojaosadati, 2017). The results obtained in this study are similar to those reported by Sahadeva et al. on the ability of *L. casei* to tolerate acidic conditions, and also the decrease in the number of them at pH 3 for 3 hours from 5.25 Log CFU/g to 3.59 Log CFU/g (Sahadeva et al., 2011). Matto et al. found that most of the bacteria studied from *Bifidobacterium animalis* species were able to tolerate pH 3, with a maximum loss of 1 Log CFU/g of bacteria (Matto et al., 2004). In a more recent study by Liu et al., *L. casei* could tolerate gastric and intestinal juices, and when exposed to gastric and intestinal juices, depending on the strain, between 72-96% of them were able to tolerate 180 minutes of exposure to the aforementioned environment (Liu et al., 2021).

Food enrichment with probiotics usually does not affect the organoleptic properties of the final product (Coman et al., 2012; Afzaal et al., 2020). The organoleptic results of the present study are similar to those reported by Sagdic et al., who observed higher scores in the color and taste of the treatment sample and thus better acceptability, but there was not much difference between the control group and the probiotic ice cream (Sagdic et al., 2012). Moreover, no significant difference in organoleptic properties of the yogurt produced from the fermentation of cashew milk using the *L. casei* was observed (Shori et al., 2022).

## 4. Conclusion

The results obtained in this study confirm the ability of the probiotic bacterium *L. casei* to grow in wheat sprouts and, consequently, transform it into probiotic wheat sprouts. The results of pH and acidity measurement, resistance to gastric juice, and TEM imaging also confirm the presence of *L. casei* in wheat sprouts.

Although no significant difference was observed between the dry matter content of the probiotic sprout and the control one, the addition of the *L. casei* not only enhanced the properties related to sprout growth significantly, but also effectively increased the total phenols and flavonoids content and its antioxidant properties. The significant increase in acidity and decrease in pH in the treated sprouts, compared with the control, also confirms the presence of *L. casei* in this study. The increase in the number of probiotic bacteria from 8.18 Log CFU/g to 11.81 ± 0.33 Log CFU/g, on the one hand, and the survival of 10.87 ± 0.28 Log CFU/g of the inoculated probiotic bacteria in the medium containing pepsin and 10.95 ± 0.12 Log CFU/g of the inoculated probiotic bacteria in the medium containing pancreatin, on the other hand, indicate that wheat sprout can be a suitable carrier for the transfer of probiotics to the human body.

The sensory acceptability of probiotic-enriched wheat sprout showed no significant difference from conventional wheat sprout. Considering the potential health benefits of probiotic sprouts, this parity in acceptability may represent a significant commercial advantage for future functional food markets.

## CRediT authorship contribution statement

**Hasan Hajjami Barkousaraei**: Conceptualization, Investigation, Methodology, Writing – original draft

**Mojtaba Mohammadzadeh Vazifeh**: Conceptualization, Methodology, Supervision, Resources, Data curation, Writing – review & editing

**Mohammad Yaghoubi-Avini**: Supervision, Data curation, Methodology, Writing – review & editing

**Golnaz Shambayati**: Methodology, Investigation, Writing – original draft

## Ethical statement

This research was conducted in accordance with appropriate ethical protocols: participation was voluntary; no coercion was used; study requirements and potential risks were disclosed to participants; informed consent was obtained; data privacy was respected; and participants retained the right to withdraw at any time.

## Acknowledgment

The authors would like to thank Arman Hedayat Noosh Alborz Food Industry Service and Research Center, Pasteur Institute of Iran, and Research Institute of Forests and Rangelands for providing technical support.

## Supporting information

S1 Table. Roots and sprouts height, means (A, C) and analysis of variance (B, D)

S 2 Table. Probiotic count (A), means (B) and analysis of variance (C)

S 3 Table. Total phenol measurement (A), means (B), analysis of variance (C)

S 4 Table. Total flavonoids measurement (A), means (B), analysis of variance (C)

S 5 Table. Antioxidant properties measurement (A), means (B), analysis of variance (C)

S 6 Table. pH measurement at minutes one and two

S 7 Table. Means (A) and Analysis of variance of pH measurement at minute one (B)

S 8 Table. Means (A) and Analysis of variance of pH measurement at minute two (B)

S 9 Table. Acidity Measurement (A), means (B) and analysis of variance (C)

S 10 Table. Dry matter measurement (A), means (B) and analysis of variance (C)

S 11 Table. Gastrointestinal juice exposure colony count (A), Means (B) and analysis of variance (C)

S 12 Table. Taste Analysis of variance

S 13 Table. Taste Means

S 14 Table. Aroma Analysis of Variance

S 15 Table. Aroma Means

S 16 Table. Texture analysis of variance

S 17 Table. Texture means

S 18 Table. Color analysis of variance

S 19 Table. Color means

S 20 Table. Appearance analysis of variance

S 21 Table. Appearance means

